# Identification of leaf rust resistance loci in hard winter wheat using genome-wide association mapping

**DOI:** 10.1101/2024.10.09.617432

**Authors:** Indira Priyadarshini Lakkakula, James A Kolmer, Rajat Sharma, Paul St. Amand, Amy Bernardo, Guihua Bai, Amir Ibrahim, Robert L. Bowden, Brett F. Carver, Jeffrey D. Boehm, Meriem Aoun

## Abstract

Leaf rust, caused by *Puccinia triticina* (*Pt*), is a serious constraint to wheat production. Developing resistant varieties is the best approach to managing this disease. Wheat leaf rust resistance (*Lr*) genes have been classified into either all-stage resistance (ASR) or adult-plant resistance (APR). The objectives of this study were to identify sources of leaf rust resistance in contemporary U.S. hard winter wheat (HWW) and to dissect the genetic basis underlying leaf rust resistance in HWW. A panel of 732 elite HWW genotypes was evaluated for response to U.S. *Pt* races at the seedling stage and at the adult plant stage in leaf rust nurseries in Oklahoma, Texas, and Kansas. Further, the panel was genotyped using Multiplex Restriction Amplicon Sequencing (MRA-Seq) and DNA markers linked to the known ASR genes *Lr18*, *Lr19*, *Lr21*, *Lr24*, *Lr37*, and *Lr42* and APR genes *Lr34*, *Lr46*, *Lr67*, *Lr68*, *Lr77*, and *Lr78*. Single nucleotide polymorphism (SNP) markers derived from MRA-Seq, DNA markers linked to the known *Lr* genes, and the phenotypic data were used for genome-wide association study (GWAS) to identify markers associated with leaf rust response. Gene postulation based on leaf rust reactions, DNA markers, and GWAS suggested the presence of *Lr1*, *Lr2a*, *Lr10*, *Lr14a*, *Lr16*, *Lr18*, *Lr19*, *Lr21*, *Lr24*, *Lr26*, *Lr34*, *Lr37, Lr39*, *Lr42*, *Lr46*, *Lr68*, *Lr77*, and *Lr78* in the HWW panel. The GWAS identified 59 SNPs significantly associated with leaf rust response, of which 20 were likely associated with novel resistance loci and can be used to enhance wheat leaf rust resistance.

## 1. INTRODUCTION

Wheat (*Triticum aestivum* L.) is one of the most important food crops globally and ranks third among U.S. field crops behind corn and soybeans (USDA Economic Research Service, 2024). In 2023, 1.8 billion bushels of wheat were harvested in the U.S. (U.S. Department of Agriculture - Economic Research Service, 2024). Winter wheat accounts for ∼70% of total U.S. wheat production. Hard winter wheat (HWW) is the most produced class of wheat and is primarily grown in the U.S. Great Plains. However, wheat production can be severely reduced due to rust diseases. Leaf rust, caused by the biotrophic fungus *Puccinia triticina* Erikss (*Pt*), is the most common wheat rust worldwide (Bolton et al., 2008). *P*. *triticina* is highly diverse for virulence to leaf rust resistance (*Lr*) genes in wheat. For instance, 30 to 60 *Pt* races are detected in the U.S. annually with virulence to multiple *Lr* genes deployed in commercial cultivars (Kolmer & Fajolu, 2022).

*Lr* genes have been classified into seedling resistance, also known as all-stage resistance (ASR), and adult-plant resistance (APR). The ASR follows the gene-for-gene model described by Flor (1955), which states that for each resistance gene in the plant, there is a corresponding avirulence gene in the pathogen. ASR is qualitative and monogenic, but it is usually race-specific and may favor the selection of virulent races of rust pathogens. Therefore, the deployment of a single ASR gene in a commercial variety usually leads to rapid loss of resistance (Bariana et al., 2006; Sucher et al., 2017). Many of the cloned ASR genes in wheat are associated with the nucleotide-binding domain and leucine-rich repeat (NLR) gene family (Feuillet et al., 2003; Huang et al., 2003; Cloutier et al., 2007; Thind et al., 2017; Yan et al., 2021; Lin et al., 2022). In contrast, wheat genotypes carrying adult plant resistance (APR) are susceptible at the seedling stage, but express resistance at the adult plant stage. APR genes can be either race-specific or non-race-specific. Non-race-specific APR genes, also called slow-rusting genes, provide partial resistance and are more durable (Krattinger et al., 2009; Moore et al., 2015), thus are desired by wheat breeding programs. As slow-rusting genes are quantitative and provide partial resistance, pyramiding multiple slow-rusting genes is needed to achieve high levels of protection.

To date, 83 *Lr* genes have been mapped in wheat and assigned gene designations (McIntosh et al., 2022; Xu et al., 2022; Kolmer et al., 2023). However, many of them are ASR genes that are either no longer effective against current *Pt* races or have not been successfully utilized in wheat breeding programs. Of the 83 characterized *Lr* genes, only eight slow rusting genes have been characterized in wheat, including *Lr34* (Krattinger et al., 2009), *Lr46* (Singh et al., 1998), *Lr67* (Hiebert et al., 2010), *Lr68* (Herrera-Foessel et al., 2012), *Lr74* (Mcintosh et al., 2016), *Lr75* (Singla et al., 2017), *Lr77* (Kolmer et al., 2018b), and *Lr78* (Kolmer et al., 2018a). To date, only *Lr34* and *Lr67* have been cloned and encode an ATP binding cassette (ABC) transporter and a hexose transporter, respectively (Krattinger et al., 2009; Moore et al., 2015). Pyramiding multiple slow rusting genes or, together with some effective ASR, is ideal for achieving durable and high levels of protection.

Kolmer and Hughes (2018) reported that different *Lr* genes were present in the hard winter wheat (HWW) grown in the southern and central Great Plains, the soft red winter wheat in the eastern states, and the hard red spring wheat in the northern Great Plains. Consequently, different *Pt* races have been found in these three major U.S. wheat growing regions (Kolmer et al., 2020). In the Great Plains, only 26% of HWW varieties grown currently were rated as highly resistant or moderately resistant to leaf rust (Onofre et al., 2023). Therefore, more varieties with durable leaf rust resistance are needed. In addition, limited information is available on genetic factors underlying leaf rust resistance in Great Plains HWW.

Advances in next-generation sequencing technologies have provided high-throughput molecular marker platforms for wheat research, which can generate thousands of genome-wide single nucleotide polymorphism (SNP) markers at an affordable cost. Genome-wide association study (GWAS) is one of the most efficient and rapid approaches to identify significant marker-trait associations (MTAs). Several GWAS have been conducted in different wheat classes to identify genomic regions associated with leaf rust resistance (Aoun et al., 2016, 2021; Sapkota et al., 2019; Fatima et al., 2020; Kaur et al., 2023). However, to date there has been no comprehensive GWAS to investigate leaf rust resistance loci/genes in contemporary HWW. The objectives of the present study were to identify sources of leaf rust resistance in contemporary HWW and to dissect the genetic basis underlying leaf rust resistance in HWW.

## 2. MATERIALS AND METHODS

### 2.1 Plant materials and genotyping

A panel of 732 HWW breeding lines and varieties was obtained from the United States Department of Agriculture – Agricultural Research Service (USDA-ARS) regional nursery program for years 2021 and 2022 (https://www.ars.usda.gov/plains-area/lincoln-ne/wheat-sorghum-and-forage-research/docs/hard-winter-wheat-regional-nursery-program/research/). These advanced wheat breeding lines (hereafter referred to as genotypes) were contributed by multiple public and private HWW breeding programs across 13 U.S. states in the Great Plains. The genotypes were submitted to the Regional Germplasm Observation Nursery (RGON), the Northern Regional Performance Nursery (NRPN), and the Southern Regional Performance Nursery (SRPN). The genotypes submitted to the NRPN and SRPN are considered elite breeding lines, and hence likely to be future variety releases for their respective breeding programs.

The panel of 732 genotypes was genotyped using Multiplex Restriction Amplicon Sequencing (MRA-Seq) (Bernardo et al., 2020) at the USDA-ARS Central Small Grains Genotyping Lab in Manhattan, KS. Single nucleotide polymorphisms (SNPs) were called using TASSEL software v.5 (Bradbury et al., 2007) and the Chinese Spring IWGSC_RefSeqv2.1 (Zhu et al., 2021) as the reference genome to assign the physical positions for each SNP markers. SNPs with missing data ≤ 65% were kept for imputation using Beagle 5 (Browning et al., 2018). Imputed markers with minor allele frequency ≥ 0.05 and heterozygosity ≤15% were used for downstream analysis. This panel was also genotyped using the DNA markers linked to known *Lr* genes *Lr18*, *Lr19*, *Lr21*, *Lr24*, *Lr34*, *Lr37*, *Lr42*, *Lr46*, *Lr67*, *Lr68*, *Lr77* and *Lr78*. Information on primer sequences and PCR protocols for these markers are available upon request from the USDA-ARS Genotyping Lab, Manhattan, KS.

### 2.2 Leaf rust evaluations at the seedling stage

Throughout multiple years, USDA-ARS coordinated evaluations of NRPN, SRPN, and RGON for multiple diseases and agronomic traits in multiple U.S. locations (https://www.ars.usda.gov/plains-area/lincoln-ne/wheat-sorghum-and-forage-research/docs/hard-winter-wheat-regional-nursery-program/research/). A total of 151 NRPN and SRPN genotypes from 2021 and 2022 were evaluated at the seedling stage against 13 U.S. *Pt* races including KFBJG, MNPSD, TCRKG, MJBJG, MCTNB, TCGJG, MHDSB, TFTSB, TNBJS, TBBGS, MBDSD, MPPSD, and TCBGS. Among these 13 races, MBDSD and MPPSD were used to evaluate only in 2022 NRPN and SRPN genotypes, whereas TCBGS was used only to evaluate 2021 NRPN and SRPN genotypes. Seedling evaluations using this large number of races, which have different avirulence/virulence phenotypes to 20 *Lr* genes in ‘Thatcher’ wheat near-isogenic lines (Table 1), facilitated the postulation of ASR genes present in NRPN and SRPN genotypes (Supplemental Table S1). Gene postulation was performed as described by Kolmer (2003). The presence of leaf rust seedling resistance genes in the NRPN and SRPN wheat genotypes was postulated based on comparing low and high IT to the IT of the *P. triticina* isolates on the Thatcher differential lines in Table 1.

**Table 1.** Virulence/avirulence phenotypes of *Puccinia triticina* races used for seedling evaluations.

| *P. triticina* races | Avirulent to leaf rust resistance (*Lr*) genes | Virulent to leaf rust resistance (*Lr*) genes |
| --- | --- | --- |
| MPPSD | *Lr2a, Lr2c, Lr11, Lr16, Lr18, Lr21, Lr28, Lr42* | *Lr1, Lr3, Lr9, Lr24, Lr26, Lr3ka, Lr17aa, Lr30, LrB, Lr10, Lr14a, Lr39* |
| MNPSD | *Lr2a, Lr2c, Lr11, Lr16, Lr18, Lr21, Lr26, Lr28, Lr42* | *Lr1, Lr3, Lr9, Lr24, Lr3ka, Lr17a, Lr30, LrB, Lr10, Lr14a, Lr39* |
| MBDSD | *Lr2a, Lr2c, Lr3ka, Lr9, Lr11, Lr16, Lr18, Lr21, Lr24, Lr26, Lr28, Lr30, Lr42* | *Lr1, Lr3, Lr17a, LrB, Lr10, Lr14a, Lr39* |
| MJBJG | *Lr2a, Lr2c, Lr3ka, Lr9, Lr11, Lr17a, Lr18, Lr21, Lr26, Lr30, Lr39, Lr42, LrB* | *Lr1, Lr3, Lr16, Lr24, Lr10, Lr14a, Lr28* |
| TNBJS | *Lr3ka, Lr11, Lr16, Lr17a, Lr18, Lr26, Lr30, Lr42, LrB* | *Lr1, Lr2a, Lr2c, Lr3, Lr9, Lr24, Lr10, Lr14a, Lr21, Lr28, Lr39* |
| MHDSB | *Lr2a, Lr2c, Lr9, Lr24, Lr3ka, Lr11, Lr30, Lr18, Lr21, Lr28, Lr39, Lr42* | *Lr1, Lr3, Lr10, Lr14a, Lr16, Lr17a, Lr26, LrB* |
| TCGJG | *Lr9, Lr16, Lr24, Lr3ka, Lr17a, Lr30, LrB, Lr18, Lr21, Lr39, Lr42* | *Lr1, Lr2a, Lr2c, Lr3, Lr26, Lr3ka, Lr11, Lr30, Lr10, Lr14a, Lr18, Lr28* |
| TCRKG | *Lr9, Lr16, Lr24, Lr17a, LrB, Lr21, Lr39, Lr42* | *Lr1, Lr2a, Lr2c, Lr3, Lr26, Lr3ka, Lr11, Lr30, Lr10, Lr14a, Lr18, Lr28* |
| TBBGS | *Lr9, Lr16, Lr24, Lr26, Lr3ka, Lr11, Lr17a, Lr30, LrB, Lr14a, Lr18, Lr42* | *Lr1, Lr2a, Lr2c, Lr3, Lr10, Lr21, Lr28, Lr39* |
| TCBGS | *Lr9, Lr16, Lr24, Lr3ka, Lr11, Lr17a, Lr30, LrB, Lr14a, Lr18, Lr42* | *Lr1, Lr2a, Lr2c, Lr3, Lr26, Lr10, Lr21, Lr28, Lr39* |
| KFBJG | *Lr1, Lr9, Lr16, Lr3ka, Lr11, Lr17a, Lr30, LrB, Lr18, Lr21, Lr39, Lr42* | *Lr2a, Lr2c, Lr3a, Lr24, Lr26, Lr10, Lr14a, Lr28* |
| TFTSB | *Lr9, Lr16, Lr18, Lr21, Lr28, Lr39, Lr42* | *Lr1, Lr2a, Lr2c, Lr3, Lr24, Lr26, Lr3ka, Lr11, Lr17a, Lr30, LrB, Lr10, Lr14a* |
| MCTNB | *Lr2a, Lr2c, Lr9, Lr16, Lr24, Lr10, Lr18, Lr21, Lr28, Lr39, Lr42* | *Lr1, Lr3, Lr26, Lr3ka, Lr11, Lr17a, Lr30, LrB, Lr14a* |

Due to seed limitations for genotypes from the 2021 RGON [except for 30 breeding lines from Oklahoma State University (OSU) HWW breeding program], a subset of 459 genotypes from the original collection of 732 genotypes were selected for further leaf rust seedling screenings. This subset included 325 RGON genotypes (2022 RGON + 30 OSU breeding lines from the 2021 RGON), 60 NRPN genotypes, and 74 SRPN genotypes (Supplemental Table S2). These 459 genotypes were evaluated at the seedling stage against five *P. triticina* races (i.e., MPPSD, MNPSD, MBDSD, MJBJG, and TNBJS), which are common races in the U.S. Great Plains (Supplemental Table S2). These five races were selected from the 13 *Pt* races used to evaluate the 151 NRPN and SRPN genotypes. MPPSD and MNPSD have been predominant races in Texas and Oklahoma in recent years (Kolmer & Hughes, 2018; Kolmer & Fajolu, 2022). Races MBDSD, MJBJG, and TNBJS have virulence to *Lr17a* [also virulent to *Lr37* because these two genes are closely linked (Singh et al., 2001; Xue et al., 2018)], *Lr16*, and *Lr21* respectively, which are common genes deployed in U.S Great Plains cultivars (Kolmer & Hughes, 2018).

The 459 genotypes were planted in a rust-free greenhouse in an augmented design, where five to seven seeds per genotype were planted in a single cell of 72-cell trays filled with a commercial ‘Ready-Earth’ soil mix (Sun Gro, Bellevue, WA, USA). The susceptible check variety ‘TAM110’ was also planted in each tray as a susceptible control. The plants were grown at 20°C/18°C (day/night) with a 16h photoperiod. Jack’s classic® all-purpose (20-20-20) fertilizer was applied to the seedlings according to manufacturer’s instructions once a week. The plants were then inoculated at the two-leaf stage (∼10 - 12 days after planting) with urediniospores suspended in Soltrol 170 mineral oil (Chevron Phillips Chemical Company, The Woodlands, TX) at a concentration of 0.01g/mL using an inoculator pressurized by an air pump (Aoun et al., 2016). The plants were then left to air dry for about 20 min before being placed in a humidity chamber for 16 −18 h, in the dark, at 18°C with 100% relative humidity. The inoculated plants were then transferred back to the greenhouse benches until evaluations. About 10 −12 days post-inoculation, plant infection types (ITs) were rated using a 0 – 4 scale (Stakman et al., 1962), where ‘0’ represents no visible symptoms, ‘;’ represents hypersensitive flecks, ‘1’ represents small uredinia with necrosis, ‘2’ represents small to medium-sized uredinia surrounded by chlorosis, ‘3’ represents medium-sized uredinia with or without chlorosis, and ‘4’ represents large uredinia without necrosis or chlorosis (Roelfs et al., 1992). Variations in uredinia size relative to the standard IT were indicated with ‘−’ and ‘+’. ITs <3 were considered resistant, whereas ITs ≥ 3 were considered susceptible. Then the 0 – 4 scale was converted to a linearized 0 – 9 scale for GWAS as described by Zhang et al. (2014) in which ITs of 0 – 3 were classified as resistant, 4 – 6 were classified as moderately resistant and ITs of 7 – 9 were classified as susceptible.

### 2.3 Leaf rust evaluations at the adult plant stage

The 459 HWW genotypes were evaluated at the adult plant stage in the field at the OSU Entomology and Plant Pathology Farm in Stillwater, Oklahoma (GPS coordinates: 36.125118, - 97.103563) in 2023 and 2024. The wheat genotypes were planted in the fall of 2022 and 2023 in 1.5 m row plots and in an augmented design. The susceptible checks (‘OK Bullet’, ‘TAM 107’, and ‘Mattern’) were planted every 50 genotypes. OK Bullet was also planted as a spreader to enhance disease pressure in the field. Spreader rows were inoculated with a bulk of 2021-2023 *Pt* isolates collected in Stillwater, Oklahoma wheat fields by spraying urediniospores suspended in Soltrol 170 three times at 10-days intervals around tillering to the jointing stage (Feekes stage 5 to 6) using a hand-held sprayer.

In Oklahoma, leaf rust severity and infection responses on flag leaves were recorded around the milk stage (Feekes stage 11.1). In 2024, disease severity and infection responses on flag leaves were recorded twice in Oklahoma (OK24-S1 and OK24-S2), with a seven-day interval. Disease severity (scale of 0-100%) corresponds to the percentage of flag leaf area covered with leaf rust pustules (uredinia) following the modified Cobb scale (Peterson et al., 1948). Infection responses were recorded as immune = no visible symptoms; R (resistant) = visible chlorosis or necrosis with no uredinia; MR (moderately resistant) = small uredinia surrounded by either chlorosis or necrosis; MS (moderately susceptible) = medium-sized uredinia without necrosis but may be associated with chlorosis; S (susceptible) = large uredinia with no chlorosis or necrosis (Roelfs et al., 1992; McIntosh et al., 1995). A combination of any two infection response categories can occur on the same leaf, with the most predominant infection response recorded first followed by the least predominant infection response. For further analysis, disease severity and infection response were combined into a single value known as the coefficient of infection (COI), which is the product of disease severity and a constant for infection response where immune = 0.0, R = 0.2, RMR = 0.3, MR = 0.4, MRMS = 0.5, MSMR = 0.6, MS = 0.8, MSS = 0.9, S = 1 (Yu et al., 2011; Aoun et al., 2016) (Supplemental Table S3).

The 732 genotypes from 2021 and 2022 NRPN, SRPN, and RGON were evaluated in field plots in Castroville, Texas in 2021 and 2022, respectively. NRPN and SRPN genotypes were evaluated in two replications based on disease severity and infection response (similar to the leaf rust rating described in Oklahoma), whereas the RGON genotypes were evaluated in an augmented design with six checks included after every 44 genotypes and based on an infection type scale of 0-9. The 0-9 scale used at the adult plant stage is described in Supplemental Table S4. Plants with IT = 0 were immune (no symptoms), plants with IT 1 – 3 were resistant, plants with IT 4 – 6 moderately resistant to leaf rust, and plants with IT 7 – 9 were susceptible. Further, 2021 NRPN, SRPN, and RGON genotypes were evaluated at the adult plant stage in 1.5 m field plots in Manhattan, Kansas in 2021 using an augmented design with three checks repeated after every 44 genotypes and scored using scale of 0-9 for infection type and 0-100% for disease severity on the flag leaf. TAM 107 and Mattern were included with the susceptible checks in Kansas and Texas. In 2022, leaf rust pressure was low in Kansas due to extreme heat conditions that suppressed the disease development. Thus leaf rust data were not recorded. Field evaluation data in Kansas and Texas is publicly available at https://www.ars.usda.gov/plains-area/lincoln-ne/wheat-sorghum-and-forage-research/docs/hard-winter-wheat-regional-nursery-program/research/ and in Supplemental Table S3.

### 2.4 Principal component analysis and linkage disequilibrium

To examine the population structure in the collection of 459 winter wheat genotypes, a principal component analysis (PCA) was performed using the filtered and imputed SNP markers and with the ‘prcomp’ function in R. The population structure was then visualized using the first two principal components. The linkage disequilibrium (LD) between all SNP marker pairs was estimated in TASSEL version 5.2 (Bradbury et al., 2007) as the square of the correlation coefficient (r^2^). To visualize the LD across the genome and each sub-genome (A, B, and D), syntenic *r^2^* values (between markers on the same chromosome) were plotted against the physical positions of SNPs in million base pair (Mb) based on the Chinese Spring reference IWGSC RefSeq. v2.1 (Zhu et al., 2021). A critical value of *r*^2^ was calculated based on the distribution of unlinked *r*^2^ (between markers on different chromosomes) that were significant at the 99% level of confidence (*P* ≤ 0.01). The parametric 95^th^ percentile of the distribution of unlinked *r*^2^ was considered as the population-specific critical value of *r*^2^, beyond which LD was likely to be caused by genetic linkage as described by Breseghello and Sorrells (2006). On the LD decay plot, a locally weighted estimated scatter plot smoother (LOESS) curve (Cleveland, 1979) was generated using the function ‘geom_smooth’ in R package ‘ggplot2’. The estimates of the extent of LD for the whole genome and each wheat sub-genome correspond to the intersections of the LOESS curves with the population-specific critical value of *r*^2^.

### 2.5 Genome-wide association mapping

Association mapping was performed using 9,858 filtered and imputed SNPs for the panel of 459 genotypes to identify significant SNPs associated with seedling ITs to five *Pt* races and with COI at the adult plant stage in Oklahoma in 2023 (COI-OK23) and 2024 (COI-OK24-S1 and COI-OK24-S2). For leaf rust field data at the adult plant stage in Texas and Kansas, GWAS was performed using ITs of the 2021 RGON genotypes (n = 300) in Texas (IT-TX21), ITs of the 2022 RGON genotypes (n = 327) in Texas (IT-TX22), and ITs and disease severity of the 2021 NRPN, SRPN, and RGON genotypes (n = 377) in Kansas (IT-KS21 and DS-KS21, respectively). Best linear unbiased estimates (BLUE) for ITs of 681 genotypes (IT-KS-TX-BLUE) across the three environments (i.e., IT-TX21, IT-TX22, and IT-KS21) were extracted from a linear mixed model using the R package ‘lme4’ (Vazquez et al., 2010; Bates et al., 2015), where genotype effect was considered as fixed and environment effect was considered as random. GWAS for IT-TX21 was performed using 8,250 SNPs, GWAS for IT-TX22 was performed using 9,062 SNPs, GWAS for IT-KS21 and DS-KS21 was performed using 9,125 SNPs, and GWAS for IT-KS-TX-BLUE was performed using 7,442 SNPs. In addition to SNP markers, DNA markers linked to known *Lr* genes *Lr21*, *Lr24*, *Lr34*, *Lr37*, *Lr46*, *Lr68*, and *Lr77* were used in the GWAS. Markers linked to *Lr18*, *Lr19*, *Lr42*, and *Lr67* were excluded from the GWAS, because these genes were found in very low frequencies (< 5%). The marker linked to *Lr78* was also excluded from the GWAS because of high percentage of missing data.

Marker–trait associations were identified using a single-locus mixed linear model (MLM) (Zhang et al., 2010) and multi-locus models, including fixed and random model circulating probability unification (FarmCPU) (Liu et al., 2016) and Bayesian-information and Linkage-disequilibrium Iteratively Nested Keyway (BLINK) (Huang et al., 2019) implemented in the R package Genomic Association and Prediction Integrated Tool (GAPIT) version 3.0 (Wang & Zhang, 2021). For these different GWAS models, we accounted for population structure (Q matrix) and family relatedness or kinship (K matrix) (Zhang et al., 2005; VanRaden, 2008). Population structure was based on PCA, whereas kinship was based on the identity-by-state relationship matrix. The association mapping models included the K matrix and the maximum number of tested principal components (PCs) for the Q matrix was limited to the first four PCs. The different GWAS models were compared using quantile-quantile (Q-Q) plots that visualize the deviation of observed −log10 (*P*) from the expected −log10 (*P*), thus determining the optimal number of PCs to be used as covariates in the model as well as the best GWAS models (MLM, FarmCPU, and BLINK). The ‘geom_point’ function in the R package ‘ggplot2’ (Wickham, 2016) was used to generate the Manhattan plots. Significant MTAs were claimed based on a false discovery rate ≤ 0.05 (Benjamini & Hochberg, 1995).

## 3. RESULTS

### 3.1 Leaf rust responses at the seedling stage

The 459 HWW genotypes displayed a range of ITs from immune response (IT = 0) to highly susceptible response (IT = 9) against five *Pt* races at the seedling stage (Fig. 1a; Supplemental Table S2). Depending on the races used, 43-60% of the genotypes showed susceptible reactions with IT ≥ 7. The lowest percentage of susceptibility was from the race MNPSD (43%) and the highest was from the race MBDSD (60%). There were significant positive correlations (0.41 – 0.76) between reactions to the five races (Supplemental Fig. S1). A total of 92 genotypes were highly resistant to moderately resistant to all five *Pt* races, with 16 each from the NRPN and SRPN, and 60 from the RGON (Supplemental Table S5).

**Figure 1.**
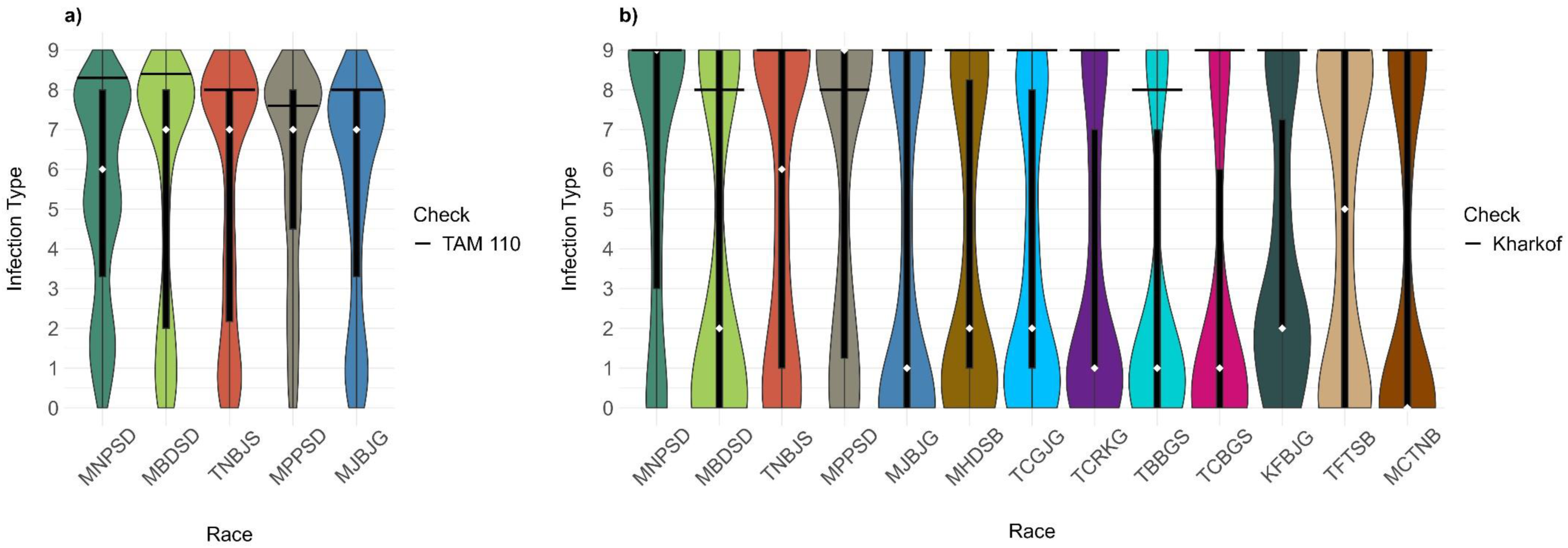
Distributions of linearized infection types for races of *Puccinia triticina* at the seedling stage in hard winter wheat genotypes. a) Distributions of infection types in 459 genotypes selected from the 2021 and 2022 NRPN, SRPN, and RGON to five *P. triticina* races. b) Distributions of infection types in 151 genotypes from the 2021 and 2022 NRPN and SRPN to 13 *P. triticina* races. Violin plots represent the density distribution of numerical observations. The box plot black rectangles represent the interquartile range, and the whiskers represent 1.5 times the interquartile range. The white diamonds represent the median. The black horizontal lines in the graphs correspond to means of the susceptible checks TAM 110 and Kharkof.

Seedling responses of 151 genotypes from the 2021 and 2022 NRPN and SRPN to 13 *Pt* races showed that the highest percentages of susceptible genotypes (44-56%) were observed for races MNPSD, MPPSD, TNBJS, and TFTSB (Fig. 1b; Supplemental Table S1). A few (n = 12) of these genotypes were treated with fungicides and thus could not be rated for infection types. A total of 13 genotypes were resistant against all 13 races, with five and eight genotypes belonging to the NRPN and SRPN, respectively. Of these 13 genotypes, eight also showed high levels of resistance across field environments (Table 2). Most of these eight genotypes carry unknown ASR genes. Based on leaf rust reactions to the 13 races, we postulated the presence of the ASR *Lr* genes *Lr1*, *Lr2a*, *Lr10*, *Lr14a*, *Lr16*, *Lr18*, *Lr21*, *Lr24*, *Lr26*, *Lr37*, and *Lr39* in the NRPN and SRPN genotypes (Supplemental Table S1). Higher frequencies of *Lr1*, *Lr16*, and *Lr26* but lower frequencies of *Lr14a*, *Lr37*, and *Lr39* were observed in NRPN genotypes than SRPN genotypes. Only a few genotypes (1– 2%) carry *Lr2a*, *L10*, *Lr11*, and *Lr18* (Fig. 2).

**Figure 2.**
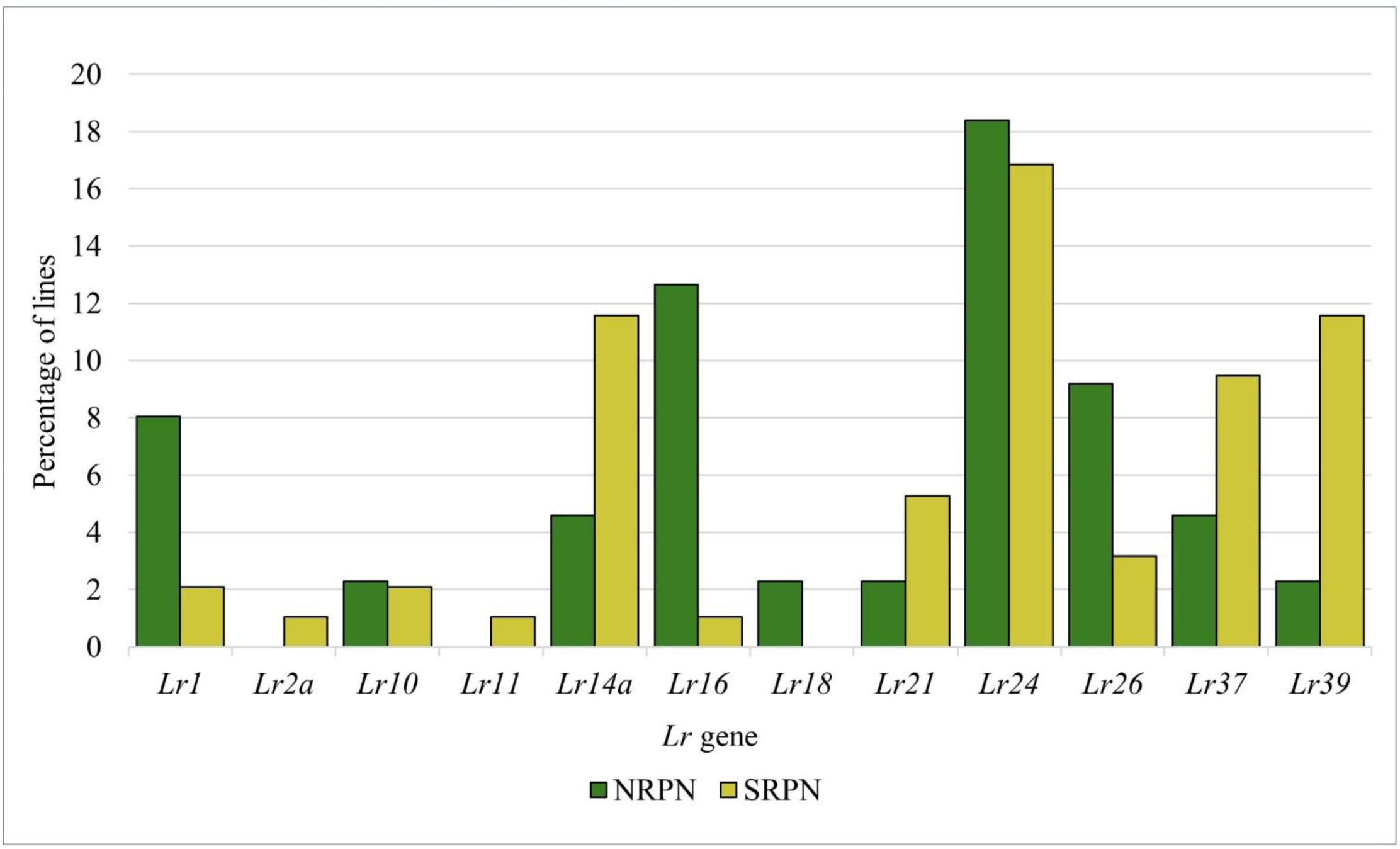
Percentages of 151 genotypes from the 2021 and 2022 NRPN and SRPN that were postulated to carry different all-stage leaf rust resistance genes based on their seedling responses to 13 *P. triticina* races.

**Table 2.**
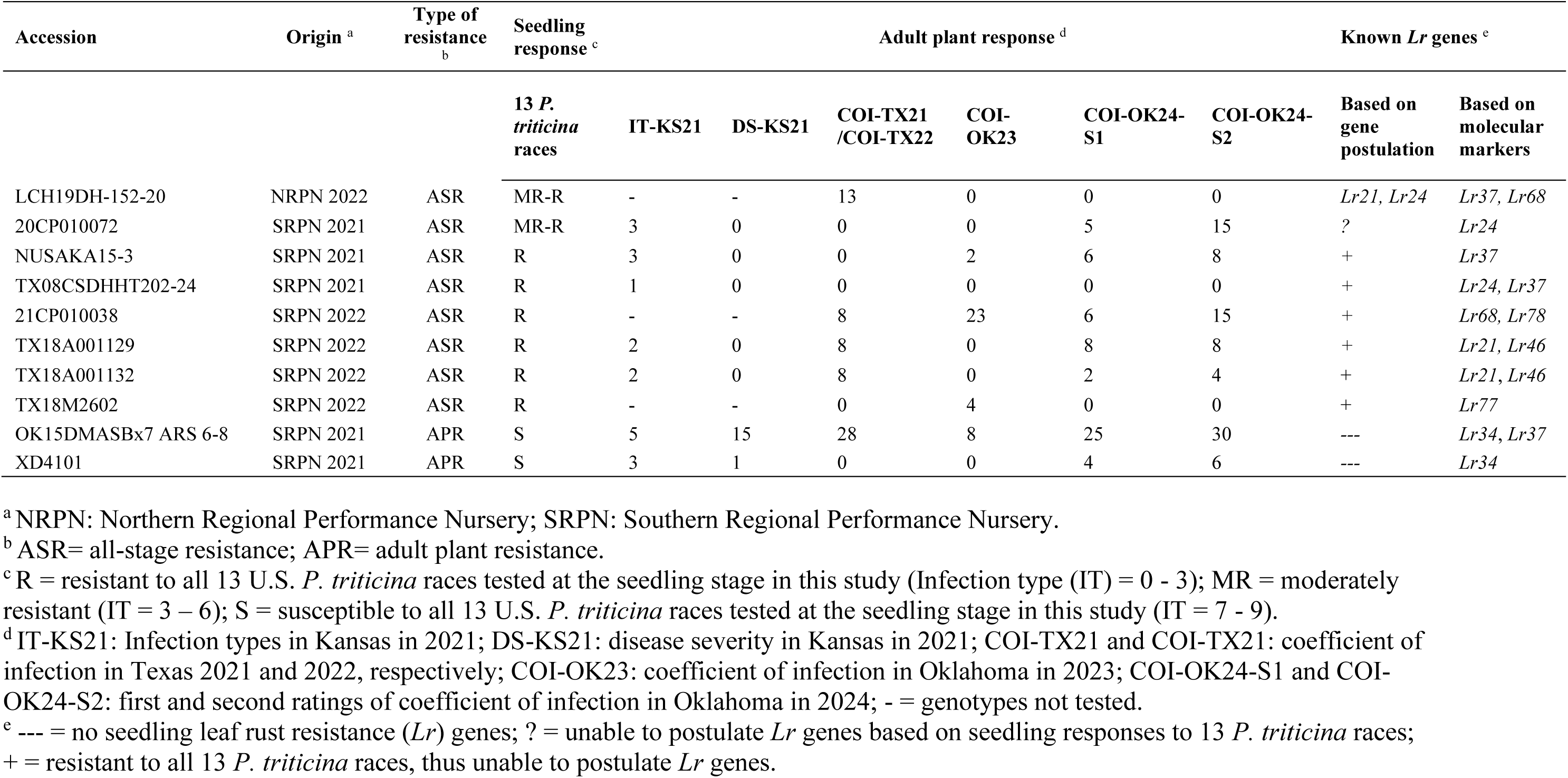
Eight sources carrying effective broad spectrum all-stage resistance and two sources carrying exclusively effective adult plant resistance among the 2021 and 2022 NRPN and SRPN genotypes.

### 3.2 Leaf rust responses at the adult plant stage

Leaf rust pressure in Oklahoma in 2023 (OK23) was moderate on susceptible checks (OK Bullet, TAM 107, and Mattern), with COI ranging from 47 to 60. Field evaluation of 459 HWW genotypes at the adult plant stage in OK23 showed that 66% (n= 301) were highly resistant (COI ≤ 20%), whereas 7% (n= 32) were susceptible (COI ≥ 60) (Fig. 3a; Supplemental Table S3). In 2024, adult plant stage evaluations in Oklahoma for the first rating (OK24-S1) showed that 53% of the genotypes (n= 240) were highly resistant, whereas 7% (n= 32) were susceptible. The subsequent rating (OK24-S2) exhibited higher disease pressure where 37% (n= 170) of the genotypes were highly resistant and 16% (n=75) were susceptible (Fig.3a; Supplemental Table S3). The susceptible checks (OK Bullet, TAM 107, and Mattern) showed COI ranging from 42 to 70 in OK24-S1 and 59 to 85 in OK24-S2.

**Figure 3.**
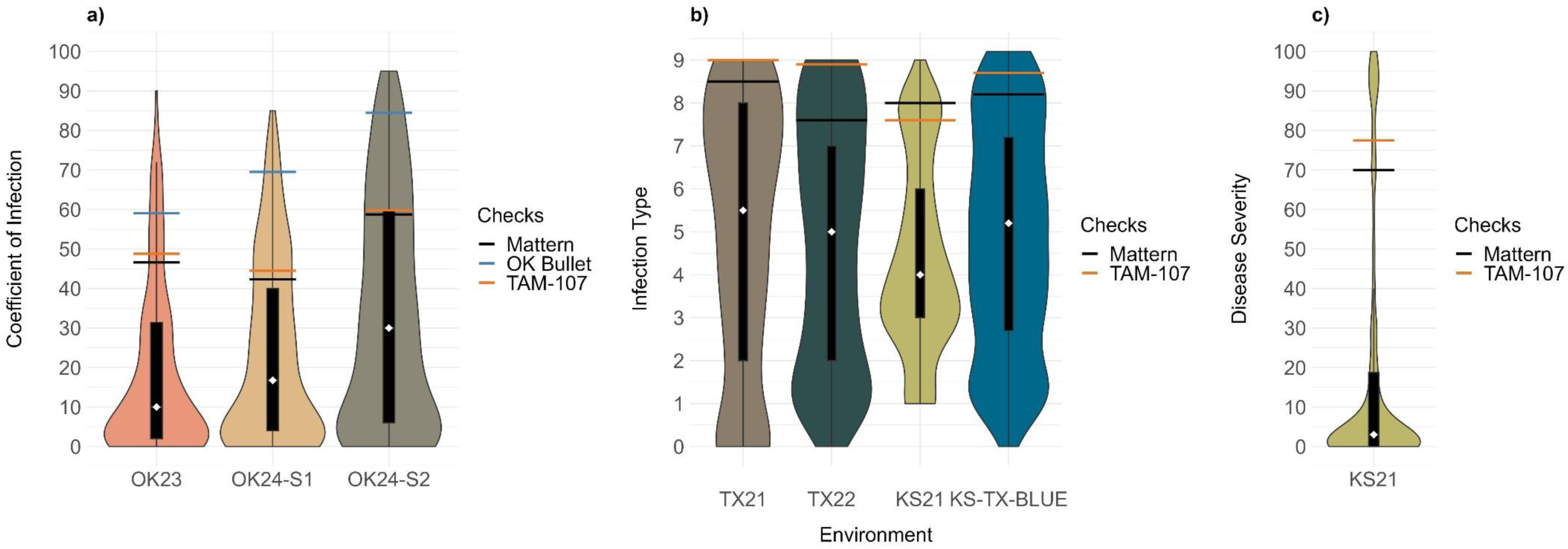
Distributions of leaf rust responses at the adult plant stage in hard winter wheat genotypes in field environments. a) Distribution of coefficient of infections of 459 genotypes selected from the 2021 and 2022 NRPN, SRPN, and RGON in Oklahoma in 2023 (OK23) and in 2024 (OK24, rated twice OK-S1 and OK-S2). b) Distributions of infection types of the 2021 RGON genotypes (n = 300) in Texas in 2021 (TX21), the 2022 RGON genotypes (n = 327) in Texas in 2022 (TX22), the 2021 NRPN, SRPN, and RGON genotypes (n = 377) in Kansas in 2021 (KS21), and best linear unbiased estimates (BLUE) of 681 genotypes across three environments, TX21, TX22, and KS21 (KS-TX-BLUE). c) Distribution of disease severities of the 2021 NRPN, SRPN, and RGON genotypes (n = 377) in KS21. Violin plots represent the density distribution of numerical observations. The box plot black rectangles represent the interquartile range, and the whiskers represent 1.5 times the interquartile range. The white diamonds represent the median. The horizontal black, blue, and orange lines in the graphs correspond to the means of the susceptible check Mattern, OK Bullet, and TAM 107, respectively.

In Texas 2021 (TX21), Texas 2022 (TX22), and Kansas 2021 (KS21), disease pressure was high with IT means ≥ 7 and disease severity means ≥ 70% for the susceptible checks TAM 107 and Mattern (Fig. 3b, c). Higher percentages of susceptible genotypes were observed in Texas compared to that in Kansas. For instance, although 80% of the genotypes (n = 294) tested in KS21 were also tested in TX21, 21% of the genotypes were susceptible (IT ≥ 7) in KS21, whereas 40% of the genotypes were susceptible in TX21 (Fig. 3b; Supplemental Table S3). Further, disease severity in KS21 for 71% of the genotypes did not exceed 20% (Fig. 3c). In TX22, 34% of the genotypes were susceptible (IT ≥ 7) (Fig. 3b). There were significant positive correlations (0.44 – 1.0) between leaf reactions in different field environments (Supplemental Fig. S1). As only few genotypes included in TX22 were also present in KS21 and TX21, there were no significant correlations between IT-TX22 and IT-TX21 and between IT-TX22 and IT-KS21 (Supplemental Fig. S2). Based on evaluations of the 2021 and 2022 NRPN and SRPN genotypes to 13 *Pt* races at the seedling stage and across field environments at the adult plant stage, we identified two lines (OK15DMASBx7 ARS 6-8 and XD4101) that carry exclusively effective APR, as they showed susceptibility at the seedling stage to all 13 races but resistance at the adult-plant stage across all field environments (Table 2).

### 3.3 Linkage disequilibrium and PCA

Of the filtered 9,858 SNPs, 5,227 (53%) were mapped to the A genome, 2,336 (24%) to the B genome, 2,204 (22%) to the D genome, and 91 (1%) were unaligned (UN) to a chromosome. SNP numbers ranged from 225 on chromosome 4B to 1,005 on chromosome 2A (Supplemental Fig. S3, S4). The highest density of SNP markers in the 1Mb window was observed on the A genome, whereas the lowest density of SNP markers was observed on the D genome. The genome-wise LD dropped to an *r^2^* threshold of 0.1 within 0.8 Mb on average (Supplemental Fig. S5). LD decayed to 0.1 at ∼ 0.9 Mb on average for genome A, to 0.7 Mb on average for genome B, and to 0.6 Mb on average for genome D (Supplemental Fig. S6). PCA using 9,858 SNPs, showed low structure in the 459 HWW genotypes, where the first two principal components (PC1 and PC2) explained 2.7% and 2.3% of genetic variation, respectively (Fig. 4). The first 10 PCs accounted cumulatively for 12.7% of the variation.

**Figure 4.**
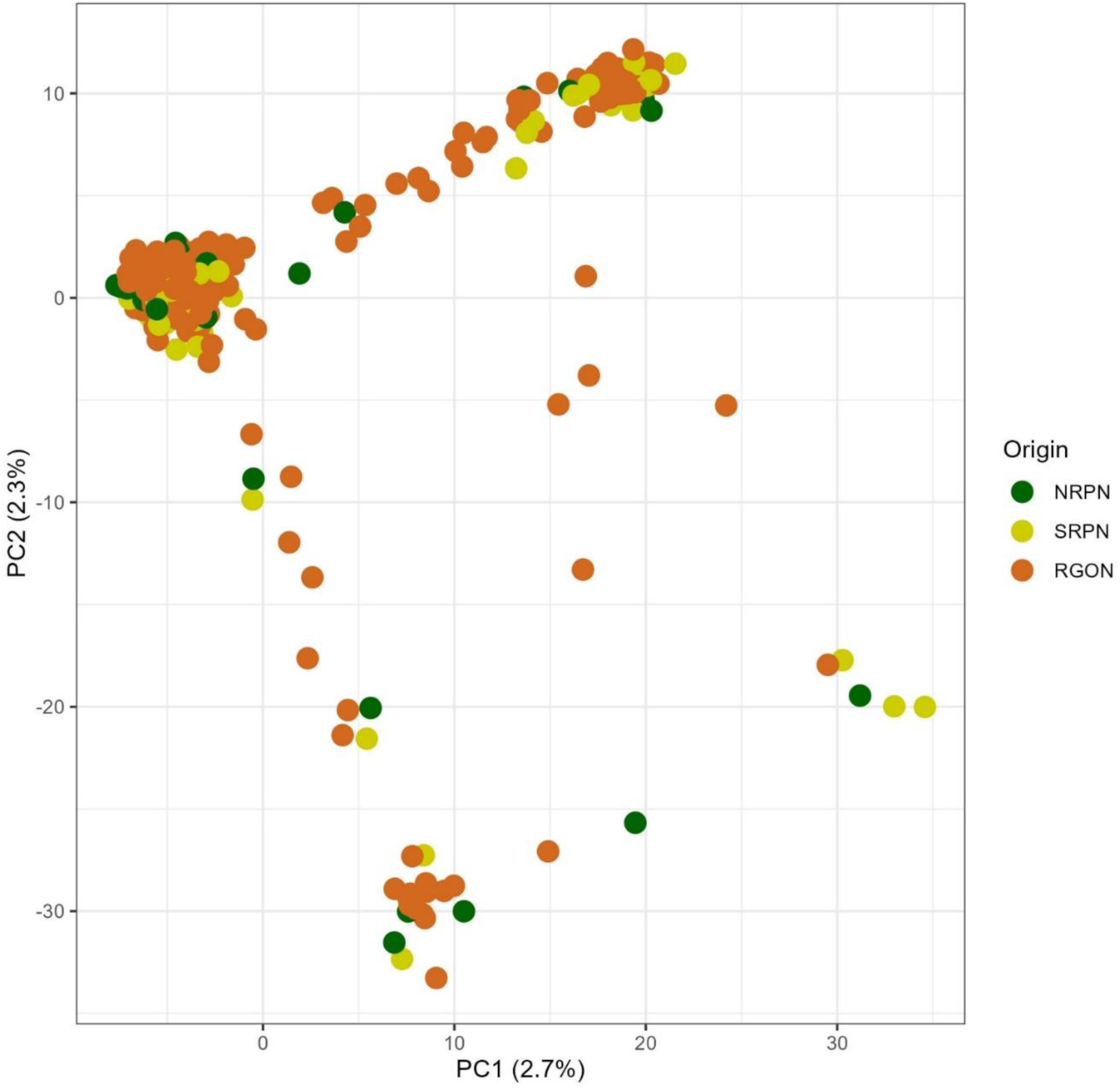
Principal component (PC) analysis obtained from 9,858 SNPs in 459 hard winter wheat genotypes selected from the 2021 and 2022 NRPN, SRPN, and RGON. The first two PCs, PC1 and PC2 explained 2.7% and 2.3% of the variation, respectively.

### 3.4. Identified *Lr* genes based on DNA markers

Based on available diagnostic markers linked to leaf rust resistance genes, the ASR genes *Lr21*, *Lr24*, and *Lr37* were found in 10%, 22% and 57% of the 459 genotypes, respectively (Fig. 5). *Lr21* is effective against races MNPSD, MPPSD, MBDSD, MJBJG, but not effective against race TNBJS. All races, except MBDSD, are virulent to *Lr24* (Table 1). Frequencies of *Lr21* and *Lr24* in the NRPN and SRPN genotypes identified by molecular markers were similar to those identified using gene postulation. In contrast to gene postulation presented in Fig. 2, *Lr37* was present in a high frequency (57%) in the NRPN and SRPN genotypes based on the molecular marker (Fig. 5). The gene postulation for *Lr37* presence is difficult because this gene has an intermediate to high IT at the seedling stage to most avirulent isolates. Thus, marker data for *Lr37* should be much more accurate than gene postulation. Although *Lr37* was present in high frequency in HWW, it is not effective against the most common U.S. *Pt.* races. The slow rusting genes *Lr34*, *Lr46*, *Lr68*, *Lr77*, and *Lr78* were found in 19%, 65%, 28%, 46%, and 15% of the genotypes, respectively (Fig. 5).

**Figure 5.**
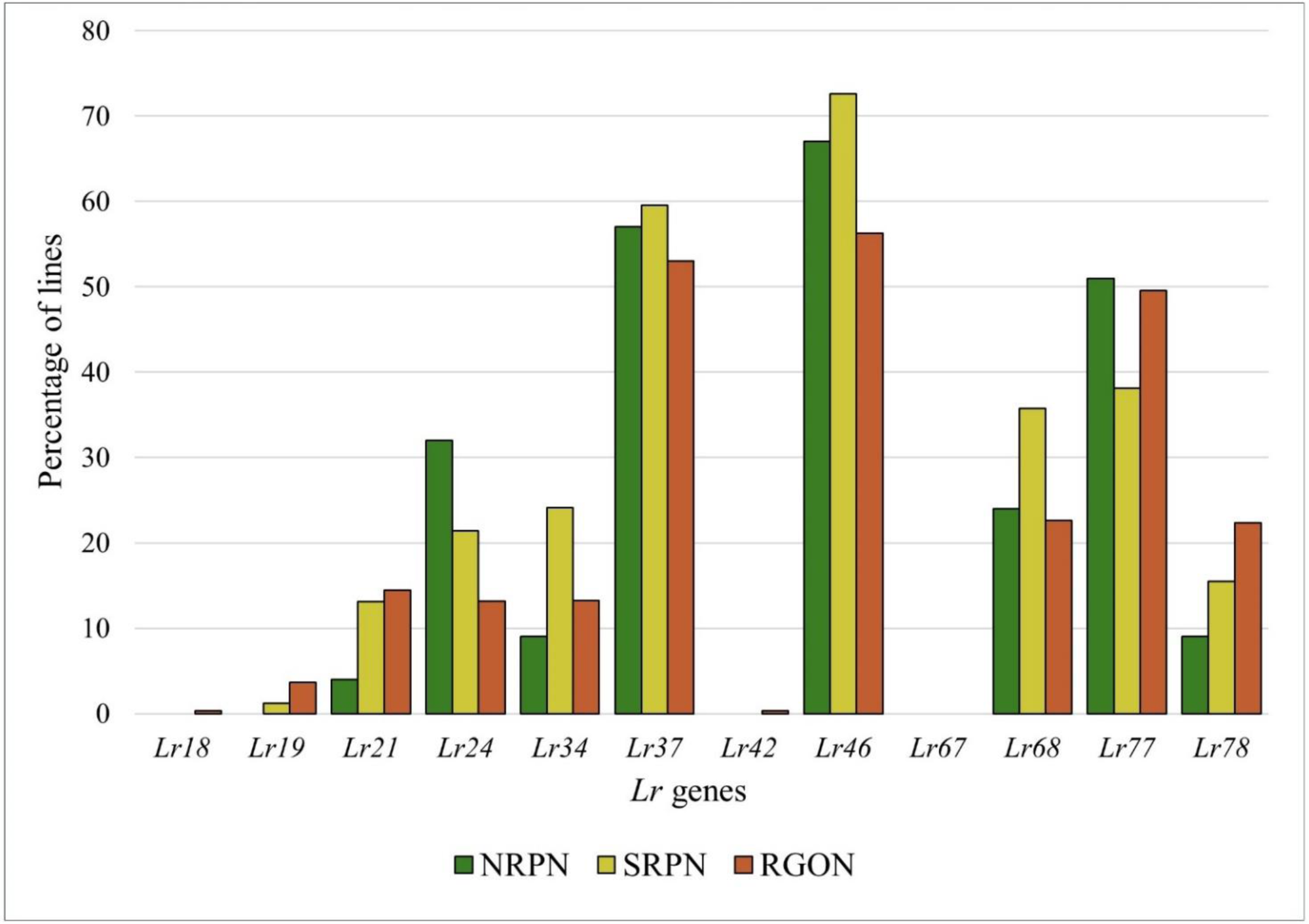
Frequencies of leaf rust resistance (*Lr*) genes in 459 hard winter wheat genotypes, selected from the 2021 and 2022 NRPN, SRPN, and RGON based on molecular markers. All stage resistance genes include *Lr18*, *Lr19*, *Lr21*, *Lr24*, *Lr37*, and *Lr42*. Slow rusting genes include *Lr34*, *Lr46*, *Lr67*, *Lr68*, *Lr77*, and *Lr78*.

### 3.5 Genome-wide association mapping

Based on Q-Q plots, BLINK outperformed MLM and FarmCPU for most traits (10 out of 13 traits). Q-Q plots of the selected BLINK model for each trait are illustrated in Fig. 6, 7. Furthermore, it was reported that BLINK has superior statistical power and computational efficiency among other models available in GAPIT 3, thus offering enhanced control over spurious associations (Huang et al., 2019; Wang & Zhang, 2021). Consequently, data from BLINK models were reported in this study (Table 3). For all traits, the K matrix was included in the GWAS models and the optimal number of PCs in the Q matrix varied based on the trait. For instance, the BLINK models for traits TNBJS, MBDSD, MJBJG, COI-OK23, COI-OK24-S1, COI-OK24-S2, IT-TX22, IT-KS21, and IT-KS-TX-BLUE included no PCs (Q matrix was not included), whereas for traits MNPSD and MPPSD, the first two PCs were included in the Q matrix. For traits DS-KS21 and IT-TX21, the first three PCs and the first four PCs were included in the Q matrix of the BLINK model, respectively.

**Figure 6.**
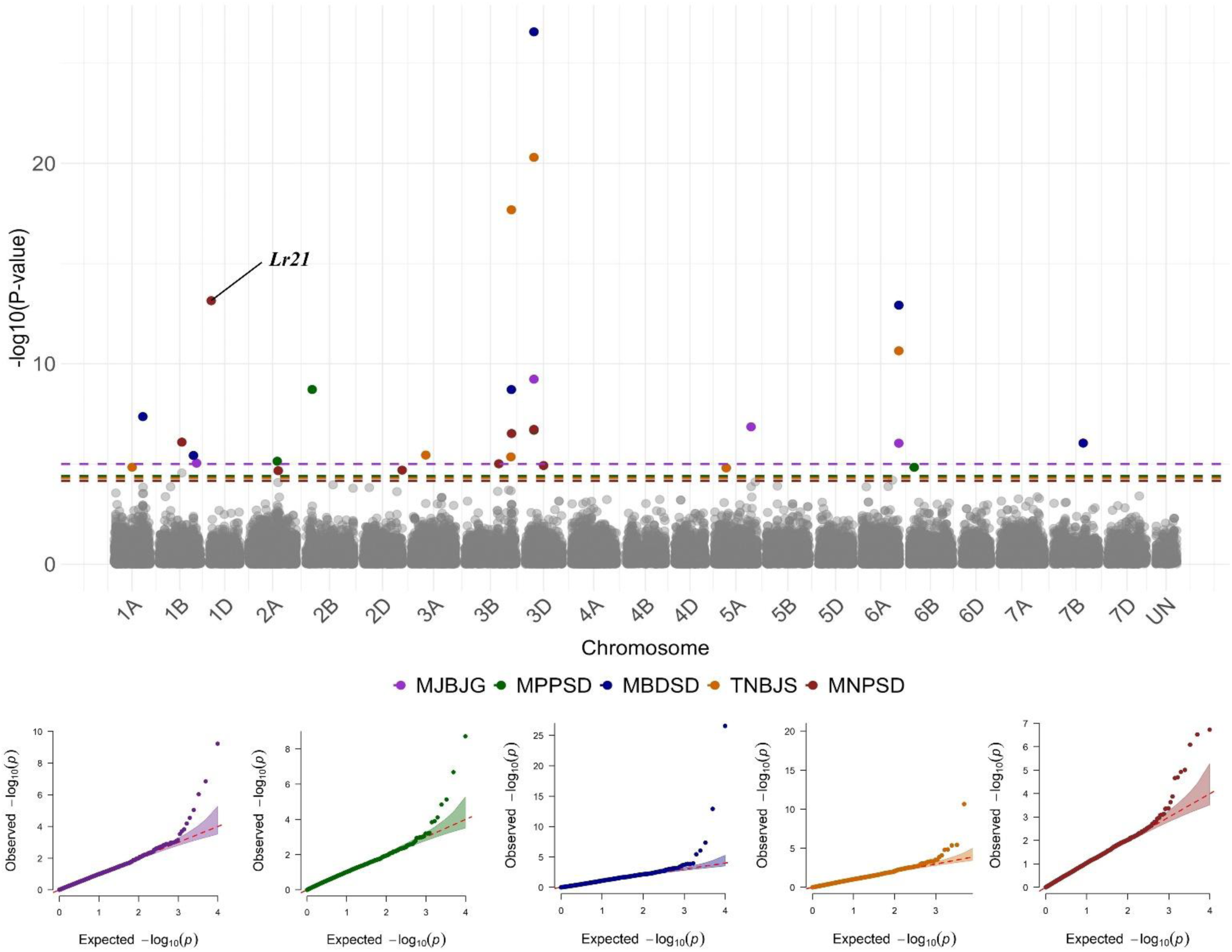
Manhattan plots showing significant SNP markers associated with leaf rust response at the seedling stage against five *Puccinia triticina* races based on the BLINK model. Colored dashed lines represent false discovery rate *P* value thresholds for respective traits. Quantile– quantile (Q-Q) plots for each trait show the expected − log10 (*P*) versus the observed − log10 (*P*) from the BLINK model.

**Table 3.**
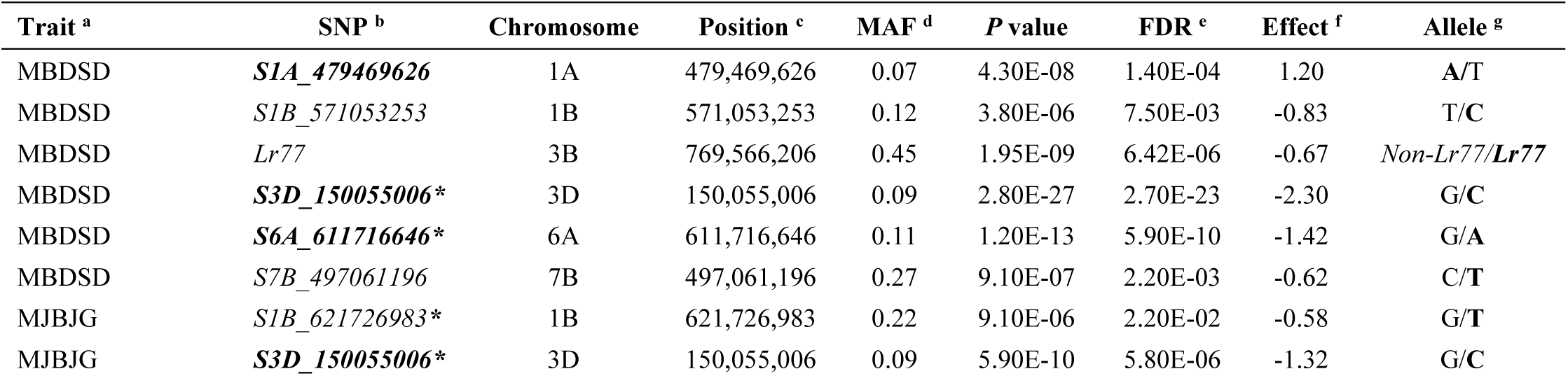
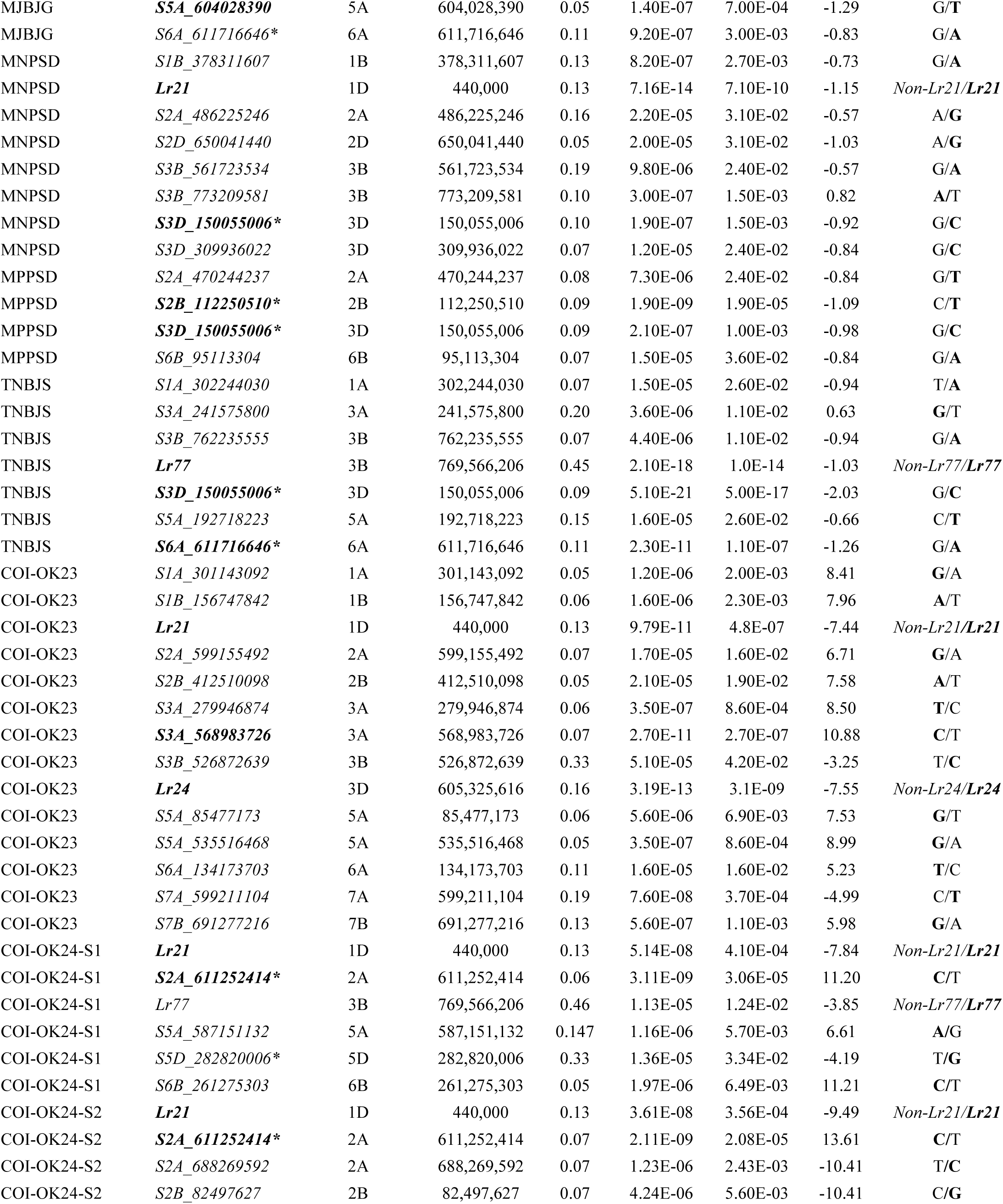
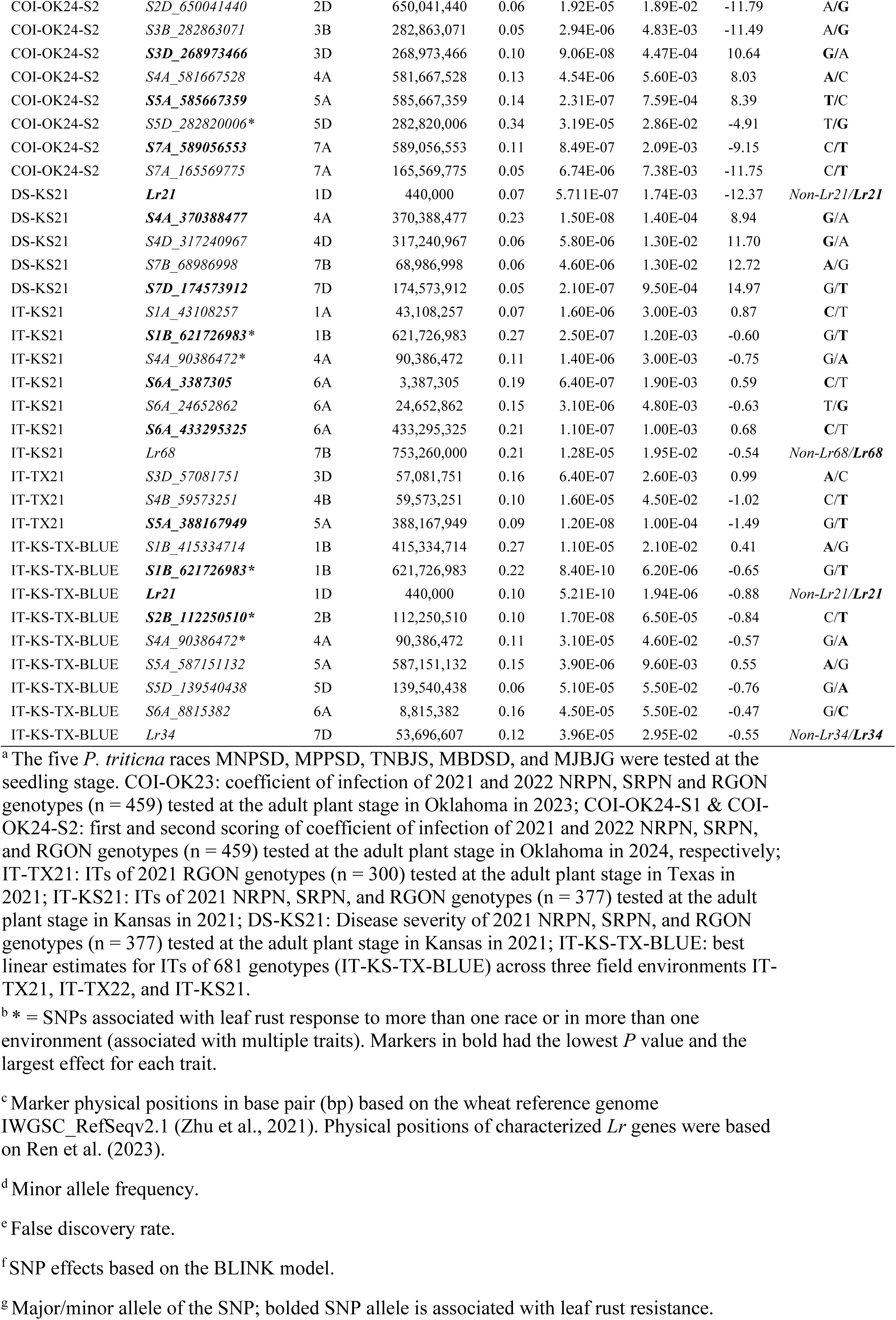
Summary of significant SNP markers associated with leaf rust responses based on the BLINK model.

Using the BLINK model, 59 SNPs (unique loci) were found to be associated with leaf rust response, of which 20 SNPs were associated with leaf rust response at the seedling stage and 42 SNPs were associated with leaf rust response at the adult plant stage. In addition, markers linked to *Lr21*, *Lr24*, *Lr34*, *Lr68*, and *Lr77* were among the significant associations. Only three markers (*S1B_621726983*, *S2B_112250510*, and *S2D_650041440*) and *Lr21* were identified at both seedling and adult plant stages. The identified MTAs at the seedling stage were distributed across 13 wheat chromosomes, whereas the MTAs identified at the adult plant stage were found on 17 wheat chromosomes (Table 3, Fig. 6, 7). Of the 732 genotypes from the 2021 and 2022 NRPN, SRPN, and RGON, 12 genotypes carried 35 – 38 favorable alleles of the 59 SNPs associated with leaf rust response in this study (Supplemental Table S6, Supplemental Fig. S7).

#### 3.5.1 Markers associated with leaf rust responses at the seedling stage

Five SNPs were associated with response to race MBDSD and were mapped to chromosomes 1A, 1B, 3D, 6A, and 7B. Markers *S1A_479469626* (i.e., SNP on chromosome 1A at 479 Mb), *S3D_150055006* and *S6A_611716646* had the lowest *P* values and had the largest effects on leaf rust response to race MBDSD (Table 3, Fig. 6). Four SNPs were associated with response to race MJBJG and were mapped to chromosomes 1B, 3D, 5A, and 6A. SNP markers *S3D_150055006* and *S5A_604028390* had the lowest *P* values and had the largest effects on response to race MJBJG. Seven significant SNPs were associated with response to race MNPSD and were found on chromosomes 1B, 2A, 2D, 3B, and 3D. *Lr21* and the SNP marker *S3D_150055006* had the lowest *P* values for leaf rust response to race MNPSD. Four SNPs were associated with response to race MPPSD and were mapped to chromosomes 2A, 2B, 3D, and 6B. Among these four SNPs, *S2B_112250510* and *S3D_150055006* had the lowest *P* values and had the largest effects on response to race MPPSD. Six SNPs were associated with response to race TNBJS and were mapped to chromosomes 1A, 3A, 3B, 3D, 5A, and 6A, of which *S3D_150055006* and *S6A_611716646* had the lowest *P* values and had the largest effects on response to race TNBJS.

The marker linked to *Lr77* was significantly associated with leaf rust seedling response to races TNBJS and MBDSD. However, *Lr77* is an APR gene; thus, an ASR gene linked to *Lr77* could be the reason for this significant association (Table 3, Fig. 6). For most of the associated markers with leaf rust response at the seedling stage (except *S1A_479469626*, *S3A_241575800*, and *S3B_773209581*), the minor alleles were associated with resistance (marker effects are negative) (Table 3). Among the 20 significant markers at the seedling stage, *S3D_150055006* and *S6A_611716646* were associated with seedling responses to five and three tested *Pt* races, respectively (Table 3, Fig. 6).

#### 3.5.2 Markers associated with leaf rust responses at the adult plant stage

Among the 42 SNPs identified at the adult plant stage, 12 SNPs were associated with COI-OK23 and mapped to chromosomes 1A, 1B, 2A, 2B, 3A, 3B, 5A, 6A, 7A, and 7B. Among these SNPs, *S3A_568983726* had the lowest *P* value and had the largest effect on COI-OK23. Markers linked to *Lr21* and *Lr24* were also among the significant associations for COI-OK23. Although, the marker linked to *Lr34* had an FDR = 0.38, it was among the most significant SNPs associated with COI-OK23 (*P* = 1.35E-3) (Table 3, Fig. 7). Four SNPs were associated with COI-OK24-S1 and mapped to chromosomes 2A, 5A, 5D, and 6B. The marker linked to *Lr77* was among the significant associations for COI-OK24-S1. Eleven SNPs were associated with COI-OK24-S2 and mapped to chromosomes 2A, 2B, 2D, 3B, 3D, 4A, 5A, 5D, and 7A. *S2A_611252414* and *S5D_282820006* were associated with both COI-OK24-S1 and COI-OK24-S2. *S2A_611252414* showed the lowest *P* values and the largest effects for both COI-OK24-S1 and COI-OK24-S2. *S3D_268973466*, *S5A_585667359*, and *S7A_589056553* were also among the markers that had the lowest *P* values and had the largest effects on COI-OK24-S2. The marker linked to *Lr21* was also among the significant associations for COI-OK24-S1 and COI-OK24-S2. Although, the marker linked to *Lr77* had an FDR = 0.29, it was among the most significant SNPs associated with COI-OK24-S2 (*P* = 4.2E-4). Four SNPs were associated with DS-KS21 and mapped to chromosomes 4A, 4D, and 7B, of which *S4A_370388477* and *S7D_174573912* showed the lowest *P* values and/or the largest effects. The marker linked to *Lr21* was also significantly associated with DS-KS21. Six SNPs were associated with IT-KS21 and were mapped to chromosomes 1A, 1B, 4A, and 6A. Of them, SNP markers *S1B_621726983*, *S6A_3387305*, and *S6A_433295325* exhibited the lowest *P* values. The marker linked to *Lr68* was also significantly associated with IT-KS21. Three SNPs, mapped to chromosomes 3D, 4B, and 5A, were associated with IT-TX21, of which *S5A_388167949* had the lowest *P* value and had the largest effect on IT-TX21. Seven SNPs were associated with IT-KS-TX-BLUE and mapped to chromosomes 1B, 2B, 4A, 5A, 5D, and 6A. *S1B_621726983* and *S2B_112250510* exhibited the lowest *P* values. *S1B_621726983* and *S4A_90386472* were associated with IT-KS-TX-BLUE and IT-KS21. Furthermore, SNP markers *S1B_621726983* and *S2B_112250510* were associated with IT-KS-TX-BLUE and also associated with seedling response to races MJBJG and MPPSD, respectively. Markers linked to *Lr21* and *Lr34* were also significantly associated with IT-KS-TX-BLUE (Table 3, Fig. 7).

**Figure 7.**
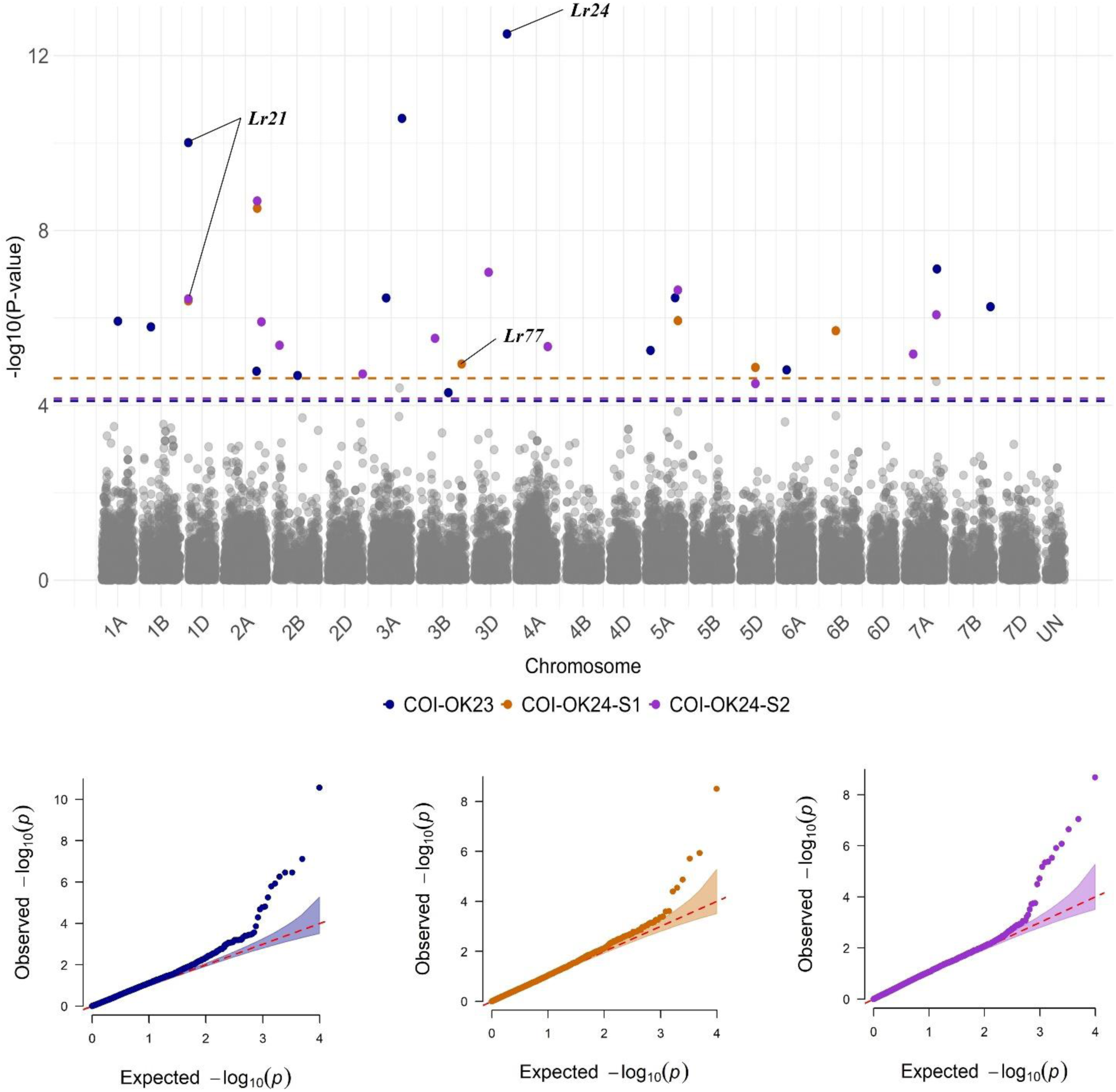
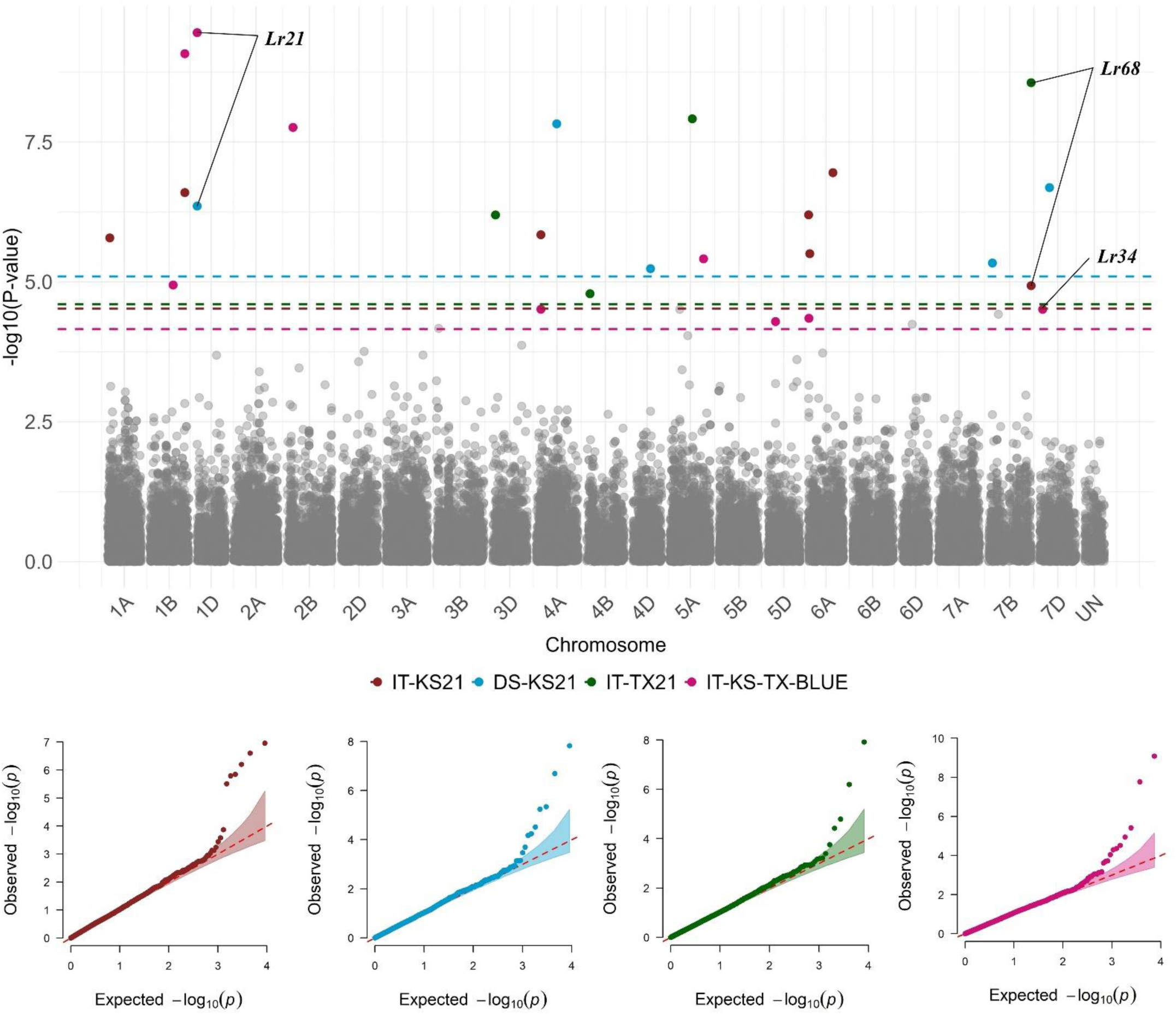
Manhattan plots showing significant SNP markers associated with leaf rust response at the adult-plant stage in different field environments based on the BLINK model. COI-OK23: coefficient of infection at the adult plant stage in Oklahoma in 2023. COI-OK24-S1 & COI-OK24-S2: first and second ratings of coefficient of infections at the adult plant stage in Oklahoma in 2024. IT-KS21: infection types at the adult plant stage in Kansas in 2021. DS-KS21: disease severity at the adult plant stage in Kansas in 2021. IT-TX21: infection types at the adult plant stage in Texas in 2021. IT-KS-TX-BLUE: best linear estimates of infection types across three field environments in Texas 2021 and 2021 and Kansas 2021. Colored dashed lines represent false discovery rate *P* value thresholds for respective traits. Quantile– quantile (Q-Q) plots for each trait show the expected − log10 (*P*) versus the observed − log10 (*P*) from the BLINK model.

## 4. DISCUSSION

This study significantly improves our understanding of the genetic basis of leaf rust resistance in the U.S. Great Plains HWW. We investigated *Lr* genes/loci in contemporary HWW using gene postulation, diagnostic molecular markers, and GWAS. Using gene postulation and diagnostic markers, HWW was found to carry 13 previously characterized ASR genes (*Lr1*, *Lr2a*, *Lr10*, *Lr14a*, *Lr16*, *Lr18*, *Lr19*, *Lr21*, *Lr24*, *Lr26*, *Lr37*, *Lr39*, and *Lr42*) and five APR slow rusting genes (*Lr34*, *Lr46*, *Lr68*, *Lr77*, and *Lr78*). Virulence to all these ASR genes, except *Lr42* has been reported in the U.S. (Kolmer, 2019; Lin et al., 2022). However, based on molecular marker data, *Lr42* was present at a very low frequency (0.3%) in the HWW germplasm, thus it was not identified in our GWAS. Molecular markers linked to the ASR genes *Lr21* and *Lr24* and the APR slow rusting genes *Lr34*, *Lr68*, *Lr77* were also among the significant associations in this GWAS study. Based on LD between markers, there were no SNP markers in strong LD (r^2^ ≥ 0.1) with DNA markers linked to known *Lr* genes (except for *Lr37*) that were used in this study. This suggests that higher-density SNP genotyping platforms may improve the discovery of additional *Lr* genes/loci in this wheat panel.

Among the 2021 and 2022 NRPN and SRPN genotypes, we identified eight sources of broad spectrum ASR originating from different HWW breeding programs in the Great Plains that were resistant to 13 *P. triticina* races and in multiple field environments. Results from molecular markers and gene postulation demonstrated that some of these eight genotypes carry ASR *Lr* genes such as *Lr21*, *Lr24*, and *Lr37*. However, virulence against these ASR *Lr* genes was present in some of the 13 races in this study. Hence these resistant sources may carry other uncharacterized effective ASR genes. Effective ASR loci were identified in this study using GWAS and further genetic studies to characterize the broad-spectrum ASR resistance genes in these eight genotypes are warranted. Among NRPN and SRPN genotypes, OK15DMASBx7 ARS 6-8, which was released as ‘Firebox’ by the OSU wheat breeding program in 2023, and XD4101 carried APR gene (s), including *Lr34* as indicated by molecular markers. However, *Lr34* provides only partial resistance (Krattinger et al., 2009), thus other unknown APR gene (s) present in Firebox and XD4101 may explain their high levels of resistance against leaf rust across multiple field environments.

Based on marker physical positions on the Chinese Spring wheat reference genome IWGSC_RefSeqv2.1 (Zhu et al., 2021), we identified previously characterized *Lr* genes/quantitative trait loci (QTLs) within genomic regions (≤ 15 Mb) of significant SNPs associated with leaf rust response in this study. Resistance loci tagged with significant SNPs were deemed novel if located in genomic regions where no previously characterized *Lr* genes (McIntosh et al., 2022; Xu et al., 2022; Kolmer et al., 2023) and QTLs were reported in 17 previous GWAS studies (Kertho et al., 2015; Aoun et al., 2016; Gao et al., 2016; Li et al., 2016; Kankwatsa et al., 2017; Pasam et al., 2017; Turner et al., 2017; Fatima et al., 2020; Kumar et al., 2020; Leonova et al., 2020; Ghulam Muhu-Din Ahmed et al., 2021; Zhang et al., 2021; Genievskaya et al., 2022; Vikas et al., 2022; Zatybekov et al., 2022; Lhamo et al., 2023; Marone et al., 2023; Tong et al., 2024) a meta-QTL analysis study (Amo & Soriano, 2022) and eight recent QTL mapping studies (Rollar et al., 2021; Xu et al., 2021; Ciechanowska et al., 2022; Wang et al., 2022; Bokore et al., 2023; Kokhmetova et al., 2023; Zhou et al., 2023; Gao et al., 2024). Five significant SNPs from the present GWAS study were found within genomic regions of previously characterized *Lr* genes (Supplemental Table S7). The significant SNP, *S1B_156747842* was found proximal to *Lr26* (Mago et al., 2002; Zhou et al., 2014). However, 94% of this HWW panel carried the resistance allele (A) of *S1B_156747842*, whereas *Lr26* on the 1RS rye translocation (Mago et al., 2002) is present in much lower frequency in this germplasm (∼13%) based on 1BL:1RS marker for the rye translocation. Therefore, *S1B_156747842* is unlikely to be associated with *Lr26*. *S6A_611716646* was found proximal to *Lr64* (Kolmer et al., 2019), however *Lr64* was recently introgressed from *Triticum turgidum* ssp. *dicoccoides* (Kolmer et al., 2019) and is probably not yet present in HWW germplasm. Likewise, the SNPs *S6A_3387305*, *S6A_8815382*, and *S6A_24652862* were detected within the genomic region of *Lr62* (Marais et al., 2009; Somo et al., 2017). *Lr62* was transferred from *Aegilops neglecta* Req. ex Bertol. to common wheat; thus, it is likely absent or present at a frequency too low in the HWW panel to be detected in our GWAS. The significant SNP *S2B_82497627*, which was associated with leaf rust response at the adult plant stage in Oklahoma was found proximal to the APR gene *Lr48* (Bansal et al., 2008). Further studies to validate the presence of *Lr48* in HWW are warranted.

Comparative mapping to previous GWAS and QTL mapping studies for leaf rust resistance, showed that 39 SNPs were co-localized within genomic regions of previously identified leaf rust resistance loci, whereas 20 were in genomic regions not known to carry previously identified leaf rust resistance loci (Supplemental Table S7). Of the 20 SNPs associated with leaf rust response at the seedling stage, 13 could be associated with novel resistance loci, including *S1A_479469626*, *S1A_302244030*, *S1B_571053253*, *S1B_37831160*7, *S2A_486225246*, *S2A_470244237*, *S3A_241575800*, *S3B_773209581*, *S3D_309936022*, *S5A_604028390*, *S5A_192718223*, *S6B_95113304*, and *S7B_497061196*. Of the 42 SNPs associated with leaf rust response at the adult plant stage, seven SNPs were possibly associated with novel resistance loci, including *S1A_301143092*, *S3A_279946874*, *S3B_526872639*, *S4A_370388477*, *S5A_388167949*, *S5D_139540438*, and *S5D_282820006*. While 39 of our identified SNPs were located within known regions carrying *Lr* genes/QTL, the discovery of 20 SNPs, which are likely associated with novel resistance loci, should enhance breeding efforts for leaf rust resistance in wheat and further resistant sources. This study analyzed identified significant loci and investigated favorable alleles associated with leaf rust resistance for potential application in breeding. There were 12 genotypes that carry 35 – 38 favorable alleles of the 59 significant SNPs from this GWAS in addition to other characterized *Lr* genes. If released as wheat cultivars, these genotypes should provide improved leaf rust resistance to rapidly evolving *P. triticina* races.

## CONCLUSION

This study provides a better understanding of the genetics underlying leaf rust resistance in hard winter wheat. Sources of ASR and APR resistance with broad effectiveness have been identified in contemporary elite hard winter wheat. Onofre et al. (2023) reported that only 26% of HWW varieties grown currently in the U.S. Great Plains are highly resistant or moderately resistant to leaf rust. Therefore, the resistant sources identified in this study can be used to enhance leaf rust resistance in HWW. Using gene postulation, molecular markers, and GWAS, we confirmed the presence of the characterized ASR genes *Lr1*, *Lr2a*, *Lr10*, *Lr14a*, *Lr16*, *Lr18*, *Lr19*, *Lr21*, *Lr24*, *Lr26*, *Lr37*, *Lr39*, and *Lr42* and the APR slow rusting genes *Lr34*, *Lr46*, *Lr68*, *Lr77*, and *Lr78*. Furthermore, we identified 59 SNPs associated with leaf rust response, of which 20 were likely associated with novel resistance loci and can be used to diversify and accelerate the deployment of resistance sources. Finally, we identified wheat genotypes with a high number of alleles conferring resistance to leaf rust that could serve as useful germplasm resources in breeding for durable leaf rust resistance.

## ACKNOWLEDGMENTS AND DISCLAIMER

This project was funded by the United States Department of Agriculture National Institute of Food and Agriculture Grant # 2023-67014-39298. The mention of trade names or commercial products in this publication is solely to provide specific information and does not imply recommendation or endorsement by the United States Department of Agriculture. The USDA is an equal opportunity provider and employer.

## DATA AVAILABILITY STATEMENT

All data generated or analyzed during this study are included in this published article and its supplementary information files. The MRA-Seq SNP data for 732 hard winter wheat genotypes are available at figshare.com/s/1dfddedeb3338844eba9

Supplemental tables are available at https://figshare.com/articles/dataset/Supplemental_Tables_sup_Identification_of_leaf_rust_resistance_loci_in_hard_winter_wheat_using_genome-wide_association_mapping_sup_/26963707?file=49066054

Supplemental figures are available at https://figshare.com/articles/figure/Supplemental_Figures_Identification_of_leaf_rust_resistance_loci_in_hard_winter_wheat_using_genome-wide_association_mapping/26963716?file=49066063

## SUPPLEMENTAL MATERIAL

### Supplemental Tables

**Supplemental Table S1**. Leaf rust infection types (IT) of 151 genotypes from the 2021 and 2022 NRPN and SRPN against 13 *P. triticina* races and possible leaf rust resistance genes in each genotype.

**Supplemental Table S2**. Leaf rust infection types (IT) of 459 hard winter wheat genotypes selected from the 2021 and 2022 NRPN, SRPN, and RGON against five *P. triticina* races.

**Supplemental Table S3**. Leaf rust responses of 732 hard winter wheat genotypes from the 2021 and 2022 NRPN, SRPN, and RGON at the adult plant stage in different field environments and *Lr* genes in each genotype based on molecular markers.

**Supplemental Table S4**. Leaf rust infection types (scale 0-9) used to rate wheat genotypes at the adult plant stage in the field in Kansas and Texas.

**Supplemental Table S5**. List of hard winter wheat genotypes from the 2021 and 2022 NRPN, SRPN, and RGON that showed seedling resistance to five *P. triticina* races MNPSD, MPPSD, TNBJS, MBDSD, and MJBJG.

**Supplemental Table S6**. Distribution of resistant alleles of significant SNP markers associated with leaf rust responses in 732 wheat genotypes originated from the 2021 and 2022 NRPN, SRPN, and RGON.

**Supplemental Table S7**. Co-localized SNPs markers associated with leaf rust response in this study with previously identified leaf rust resistance genes/QTLs.

### Supplemental Figures

**Supplemental Figure S1.** Pearson correlations between reactions of 459 hard winter wheat genotypes to five *Puccinia triticina* isolates tested at the seedling stage.

**Supplemental Figure S2.** Pearson correlations between leaf rust reactions of hard winter wheat genotypes in field environments.

**Supplemental Figure S3.** Chromosome-wise distribution of 9,858 SNP markers in 459 hard winter wheat genotypes selected from the 2021 and 2022 NRPN, SRPN, and RGON.

**Supplemental Figure S4.** Density of SNP markers (n = 9,858 SNPs) per 1Mb window on wheat chromosomes in a set of 459 hard winter wheat genotypes selected from the 2021 and 2022 NRPN, SRPN, and RGON.

**Supplemental Figure S5.** Scatter plot showing linkage disequilibrium (LD) decay across the genome.

**Supplemental Figure S6.** Scatter plot showing linkage disequilibrium (LD) decay in sub genomes A, B, and D.

**Supplemental Figure S7.** Percentages of 732 wheat genotypes originated from the 2021 and 2022 NRPN, SRPN, and RGON carrying different number of resistant alleles of the 59 significant SNPs associated with leaf rust response.

